# Boosting GPT Models for Genomics Analysis: Generating Trusted Genetic Variant Annotations and Interpretations through RAG and fine-tuning

**DOI:** 10.1101/2024.11.12.623275

**Authors:** Shuangjia Lu, Erdal Cosgun

## Abstract

Large language models (LLMs) have acquired a remarkable level of knowledge through their initial training. However, they lack expertise in particular domains such as genomics. Variant annotation data, an important component of genomics, is crucial for interpreting and prioritizing disease-related variants among millions of variants identified by genetic sequencing. In our project, we aimed to improve LLM performance in genomics by adding variant annotation data to LLMs by retrieval-augmented generation (RAG) and fine-tuning techniques. Using RAG, we successfully integrated 190 million highly accurate variant annotations, curated from 5 major annotation datasets and tools, into GPT-4o. This integration empowers users to query specific variants and receive accurate variant annotations and interpretations supported by advanced reasoning and language understanding capabilities of LLMs. Additionally, fine-tuning GPT-4 on variant annotation data also improved model performance in some annotation fields, although the accuracy across more fields remains suboptimal. Our model significantly improved the accessibility and efficiency of the variant interpretation process by leveraging LLM capabilities. Our project also revealed that RAG outperforms fine-tuning in factual knowledge injection in terms of data volume, accuracy, and cost-effectiveness. As a pioneering study for adding genomics knowledge to LLMs, our work paves the way for developing more comprehensive and informative genomics AI systems to support clinical diagnosis and research projects, and it demonstrates the potential of LLMs in specialized domains.

## 1. Introduction

Variant annotation data provide detailed information about genetic variants and their biological and clinical implications. It contains several aspects of genetic variations, including locations, types, and effects. Annotations are curated in datasets, such as Clinvar (1) and gnomAD (2), derived from large projects, experiments, and clinical reports. In addition, variant effect prediction tools, including SnpEff (3), VEP (4), help interpret unreported variants. To interpret and prioritize variants from the millions of variants called from sequencing data, researchers and health providers need to gather variant annotations from these datasets and tools, and then analyze them based on factors such as previous records of affected genes and diseases, variant allele frequency, and predicted molecular effects. This process is time-consuming and demands substantial human resources.

Large language models (LLMs) such as GPT-4 (5) and Llama (6) have exhibited extraordinary performance in various tasks and domains, offering opportunities for improvement and automation in genomics. Previous studies have demonstrated the potential of LLM in genomics domains, including polygenic risk prediction (7), identification of causal genes in genome-wide association studies (GWAS) (8), and the screening of clinical trials (9). However, current LLMs lack genomics domain knowledge such as variant annotations, which limits their performance in genomics. By incorporating variant annotations, along with advanced language comprehension and reasoning capability, LLMs can assist disease analysis by providing annotation more efficiently, eliminating the need for searching from databases and running tools, and moreover, offering interpretations based on their capabilities and knowledge, thereby supporting a wide range of downstream tasks in this domain.

There are two common methods to incorporate domain-specific knowledge into LLMs: retrieval-augmented generation (RAG) and fine-tuning. The effectiveness and tradeoffs of these two methods are still under debate (10; 11). Fine-tuning is a continuous training process that exposes the pre-trained model to a smaller, domain-specific dataset so that the model can adjust its weights and adapt to the specific task. In this paper, we employed supervised fine-tuning, using question and answer pairs to guide the model’s weight changing towards generating our desired responses. Unlike fine-tuning, RAG does not modify the pre-trained model itself (12). Instead, RAG improves accuracy and relevance in model answer generation by including relevant external information. When using RAG, the user query is not directly inputted into the model for generation. After receiving a user query, we search the external data store to retrieve relevant information, and then construct input prompts using both the user query and the retrieved information and send this prompt to the LLM to generate an enhanced response.

To harness the power of LLMs in genomics, we integrated genomics domain knowledge, specifically 190 million variant annotations, into GPT-4o and GPT-4 models through RAG and fine-tuning, which significantly improved the model’s ability to provide accurate variant annotations and enhanced interpretations. Additionally, we compared the effectiveness of RAG and fine-tuning in injecting variant annotation data to LLMs and evaluated two methods in terms of accuracy, data volume, and cost-effectiveness. As a pioneering study in application of LLMs in genomics, our project paves the way for developing more comprehensive and powerful genomics AI tools to assist in clinical and research uses.

## 2. Methods

### Variant annotation datasets

Variant annotation data records genetic variants and their health implications, which is essential for variant interpretation and prioritization in disease diagnosis and research projects. In this study, we used 4 public variant annotation datasets: ClinVar (v.2024-06-03) (1), gnomAD (v4) (2), GWAS Catalog (v.1.0) (13), pharmGKB (14; 15), and 1 annotation tool: SnpEff (v.5.2c) (3). All annotations are based on the GRCh38 human reference genome. ClinVar contains comprehensive information on 2,897,556 clinically relevant variants and their corresponding genes and phenotypes. In gnomAD, we used the exome dataset, which records 183,717,261 variants and their allele frequencies from diverse populations. From GWAS Catalog, we downloaded all associations, which curates 625,113 variants identified by GWAS studies. For PharmGKB, we used the clinical annotation and variant annotation files, encompassing 41,287 variants and their relationships with diseases or drug responses. We then applied SnpEff to predict the functional effects of all ClinVar variants using the following command line:

~~~
java −Xmx8g − jar snp Eff.jar −v −stats
       clinvar.html GRCh38. mane.1.2. ensembl
       clinvar_20240603.noinfo.vcf >
       clinvar_20240603.noinfo.snpEff.vcf
~~~

### 2.2 Preparing prompts for fine-tuning GPT-4

Fine-tuning GPT-4 requires training and validation data to be formatted in JSON Lines (jsonl). In our study, we randomly selected 3,000 variants for the training dataset and 1,000 variants for the validation dataset from the ClinVar dataset. Variant annotations in ClinVar are stored in vcf format. To construct training and validation prompts in jsonl format, we first extracted 13 informative annotation fields (chromosome, position, reference allele, alternative allele, dbSNP ID, Variation ID, Allele ID, canonical SPDI, molecular consequence, germline review, germline status, gene, condition) for each variant using bcftools (v.1.20) (16) and then converted the extracted information into jsonl format using the awk command. Each prompt consists of a system message, a user question, and an expected assistant output. The users question involves querying the full annotations of a variant by providing either the chromosome and position or the dbSNP ID as input. Below is an example command for generating prompts, along with an example training prompt:

~~~
bcftools query clinvar20240603.vcf.gz − f “%
     CHROM\ t%POS\ t%REF\ t%ALT\ t%INFO/RS\ t%ID\ t
     %INFO/ALLELEID\ t%INFO/CLNHGVS\ t%INFO/MC\
     t%INFO/CLNSIG\ t%INFO/CLNREVSTAT\ t%INFO/
     GENEINFO\ t%INFO/CLNDN” | **awk** ‘ { split ($9,
     - a, - “|“) ; split ($12, -b, - “:“) ; -print- “{\”
     messages \ “: - [{\ “role \ “: - \” system \ “, - \”
     content \ “: - \”You - are - an - expert - genomics –
     annotator. \ “}, - {\” role \ “: - \” user \ “, - \”
     content \ “: - \”Name : chr “$1 “” $2 “| Task :
     Provide - full - annotation - information - for –
     the - given - Name - including - Gene (s), Protein
     - change, Condition (s). \ “}, - {\” role \ “: - \”
     assistant \ “, - \” content \ “: - \”GRCh38
     - Chromosome : - chr “$1 “| GRCh38 – Location : - “$2
     “|REF: - “$3 “|ALT: - “$4 “| dbSNP - ID : - r s “$5 “|
     VariationID : - “$6 “| AlleleID (s) : - “$7 “|
     Canonical - SPDI : - “$8 “| Molecular
     consequence : - “a [2 ] “| Germline -
     classification : - “$ 10 “| Germline - review
     status : - “$ 11 “| Gene : - “b [1 ] “| Condition (s) :
     -” $ 13 “\ “} ] } “} ‘ | **sed** “s/rs./nan/g” | less
     > clinvar_20240603.jsonl
{ “messages “: [{ “role “: “system “, “content “:
     “You - are - an - expert - genomics - annotator. “
     }, { “role “: “user “, “content “: “Name :
     chr1_69134 | Task : - Provide - full - annotation
     - information - for - the - given - Name -
     including - Gene (s), Protein - change,
     Condition (s). “}, { “role “: “assistant “, “
     content “: “GRCh38 - Chromosome : -chr1 |
     GRCh38 - Location : -69134 |REF: -A| ALT: -G|
     dbSNP - ID : -nan | VariationID : -2205837 |
     AlleleID(s) : -2193183 | Canonical - SPDI : -
     NC_000001.11 : g.69134A>G| Molecular -
     consequence : -missense_variant | Germline –
     classification : -Likely_benign | Germline –
     review-status : -criteria_provided,
     _single_submitter | Gene : -OR4F5 | Condition (
     s) : - not_specified “} ] }
~~~

To further understand the fine-tuning performance, we also fine-tuned the model to predict individual fields such as gene, condition, and dbSNP ID, using chromosome:position as input. In this approach, we reduced the number of fields in the assistant’s output while maintaining a similar format for the training and validation prompts.

### 2.3 Preparing data for building RAG for GPT-4o

RAG supports diverse input formats, including pdf, txt, and csv. The core of RAG is its search retrieval process. For our study, we selected the csv format to create the search index from annotations, chunking them line by line. We designed our csv file with 5 columns: chromosome:position, dbSNP ID, gene, condition, and all other annotations. This format ensures that chromosome position, dbSNP ID, gene, and condition of variants are searchable.

We integrated annotations of approximately 190 million variants from 5 datasets - ClinVar, gnomAD, SnpEff, GWAS Catalog, PharmGKB - into the search index. Specific annotation fields extracted from each dataset are detailed in Supplementary Table1. Additionally, we added the source dataset name for each variant to track the origin of the information and provided search URLs for ClinVar, gnomAD, and PharmGKB variants facilitating users in accessing more variant details from these websites. We extracted information from each dataset using bcftools and converted the information into csv format. Missing annotation fields are represented by “NA”. Below is an example command for generating RAG input and an example CSV line:

~~~
bcftools query clinvar_20240603.vcf.gz − f “%
      CHROM\ t%POS\ t%REF\ t%ALT\ t%INFO/RS\ t%ID\ t
      %INFO/ALLELEID\ t%INFO/CLNHGVS\ t%INFO/MC\
      t%INFO/CLNSIG\ t%INFO/CLNREVSTAT\ t%INFO/
      GENEINFO\ t%INFO/CLNDN” | **awk** ‘ { s p l i t ($12
     , - b, - “:“) ; - gsub (/, /, - “; “, - $ 13) ; - p r i n t - “
      chr “$1 “: “$2 “, r s “$5 “,” b [1 ] “, “$ 13 “, \ “
      GRCh38 chr : chr “$1 “\ tGRCh38 pos : “$2 “\
      treference_allele : “$3 “\
      talternative_allele : “$4 “\ tdbSNP ID : rs “$5
      “\ tVariation_ID : “$6 “\ tAllele_ID : “$7 “\
      tcanonical_ SPDI : “$8 “\
      tmolecular_consequence : “$9 “\
      tgermline_review : “$ 10 “\ tgermline_status
      : “$11 “\ tGene : “b [1] “\ tCondition : “$ 13 “\
      tsource : clinvar \ tclinvar_URL : https://www.ncbi.nlm.nih.gov/clinvar/variation /” $6
      “/ \”“} ‘ | **sed** “s/rs \. /na/g” | **sed** “s /\ “
      See_Cases\ “/See_Cases/g” >
      clinvar_20240603.5col.csv
chr1:69134, na, OR4F5, not_specified, “
      GRCh38 chr : chr1 --GRCh38 pos : 69134--
      reference_allele:A- -alternative_allele:G
      ----dbSNP_ ID:na-Variation_ID:2205837--
      Allele_ID :2193183---canonical_SPDI :
      NC_000001.11 :g.69134A>G--
      molecular_consequence :SO:0001583 |
      missense_variant---germline_review :
      Likely_benign---germline_status :
      criteria_provided, _single_submitter-Gene
      : OR4F5--Condition : not_specified-source :
      clinvar--clinvar URL : https://www.ncbi.nlm.nih.gov/clinvar/variation/2205837/ “
~~~

### 2.4 Evaluating base GPT-4o and GPT-4 performance on variant annotations

To evaluate the performance of the base GPT models, we randomly sampled 3 sets of 100 variants and extracted their corresponding genes and conditions from the ClinVar dataset as our true set. We assessed the models on predicting genes and conditions using either chromosome:position or dbSNP ID of the variants as input. One example query to identify the gene of a variant using chromosome:position is “Provide corresponding gene name for variant chr2:96799611 in GRCh38. Only reply with the gene name.” We tested the models with a temperature setting of 0. For gene prediction, we measured accuracy using the exact match between the expected gene name and the model’s output. For condition prediction evaluation, we computed the Jaro similarity between the expected condition and the model’s output, considering a similarity score greater than 0.8 as a match.

We then randomly sampled 3 sets of 100 variants from ClinVar that are located in the top 10 well-studied genes (17) to further evaluate base model performance, considering that the models are more likely to have been exposed to information about these genes during pretraining. The gene prediction accuracy was measured using the same methods as previously described.

### 2.5 Fine-tuning GPT-4 using variant annotations

After converting variant annotation vcf files into training and validation prompts in jsonl format, we fine-tuned the GPT-4 model (v.0613) on the Azure OpenAI platform using these prompts. The fine-tuning process involved 3,000 training variants and 1,000 validation variants, with a batch size of 6 over 3 epochs. We used a learning rate multiplier of 1 for the training.

### 2.6 Building RAG for GPT-4o

We built a RAG system for GPT-4o using variant annotation data on the Azure AI Search platform. The core objective of the RAG system is to build a search index that achieves optimal retrieval performance. We employed a full-text keyword search method and chunked variant annotations line by line. The workflow of building the search index consists of 3 steps: adding the data source, creating the search index schema, and loading data into the search index.

We used Azure blob storage as the data source. Due to a limitation of the Azure AI Search S1 pricing tier, which restricts file size to less than 4 million characters, we split the annotation CSV files accordingly, stored them in Azure Blob Storage, and added the storage as a data source in Azure AI Search. Next, we created a search index schema based on our CSV file structure, ensuring that the five columns - chromosome:position, dbSNP ID, gene, condition, and content (all other annotations) - were retrievable and searchable. We then loaded 190 million annotation records into the search index using an indexer. Each column’s data was processed by the Standard Lucene Analyzer to remove non-essential words and phrases. Phrases and hyphenated words were split into component parts to enhance search efficiency. The variant annotation data was then tokenized and stored in the search index.

Our search index allows for searching variants based on chromosome:position, dbSNP ID, gene, condition, and other annotation information. When a user query is received, it undergoes the same analyzer and tokenizer as indexing. Then it searches for matching documents from our annotation datasets, ranks based on relevance scores, and retrieved top results. Finally, we added the search index to GPT-4o on Azure OpenAI using the ‘Add Your Own Data’ feature.

### 2.7 Evaluating models after fine-tuning and RAG

We evaluated the performance of our models based on output accuracy. For the fine-tuned models, to measure the model’s ability to memorize the trained information and accurately reproduce the desired output, we randomly selected 100 variants from the 3,000 variants training set and counted the exact matches of each annotation field between the outputs and the true set. For the RAG models, we also randomly selected 100 variants from each of the 5 input datasets and counted the exact matches for each annotation field. The parameters we used for evaluating the RAG GPT-4o model were max tokens=1600, temperature=0, and top p=0.5.

## 3. Results

### 3.1 Evaluating base GPT-4o and GPT-4 model performance on variant annotations

We assessed the variant annotation knowledge of GPT-4o (v.2024-05-13) and GPT-4 (v.0613) by the task of predicting corresponding genes and conditions for input variants. We represented input variants using either chromosome and position or dbSNP ID in the input prompts, and instructed the models to provide gene names or conditions. Model performance was evaluated by exact matches for gene name predictions and similarity scores for condition predictions (Methods).

Both GPT-4o and GPT-4 exhibited less than 2% gene prediction accuracy when tested on 3 sets of 100 variants randomly selected from the 2.8 million variants in the ClinVar dataset (Figure 1a). Considering that the models are more likely to have been exposed to well-studied genes during their pre-training, we refined our test sets to only include variants from the 10 most well-studied genes (TP53, TNF, EGFR, VEGFA, APOE, IL6, TGFB1, MTHFR, ESR1, AKT1) (17). Under these conditions, GPT-4o achieved a prediction accuracy of 0.68, while GPT-4 achieved an accuracy of 0.487 when using chromosomes and positions of variants as input. However, accuracy remained low when using dbSNP ID of variants as inputs. For condition prediction, both models demonstrated low accuracy regardless of whether the input was chromosome and position or dbSNP ID.

**Fig. 1.**
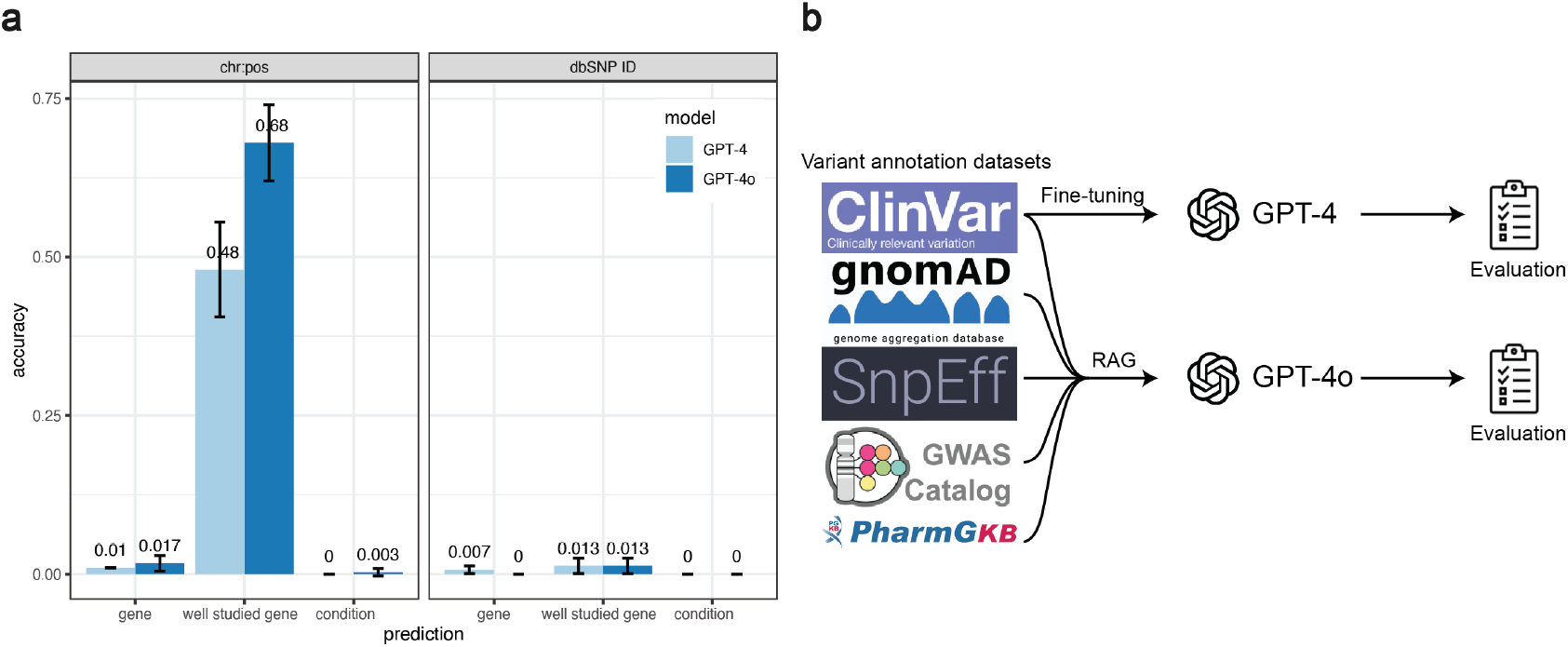
Base GPT models have limited variant annotation knowledge. (a) Performance of the base GPT-4 and GPT-4o models in predicting corresponding genes and conditions from user input variants represented in chromosome:position (chr:pos) or dbSNP ID formats. (b) Overview of our approach to improve GPT model performance in variant annotations. We integrated 5 variant annotation datasets into GPT models through RAG and fine-tuning, then evaluated their improvements.

Our results indicated that while the base GPT models show improved performance in predicting well-studied genes, their overall knowledge of variant annotation is very limited. To address this limitation, we incorporated variant annotation data into these models using RAG and fine-tuning. We then evaluated the improvements after adding this domain-specific knowledge (Figure 1b).

### 3.2 Incorporating variant annotations into GPT-4o through RAG

We collected clinically relevant variant annotations from 4 datasets: ClinVar (1), gnomAD (2), GWAS Catalog (13), and PharmGKB (14; 15) and 1 tool: using SnpEff (3) to predict functional the effect of ClinVar variants (Supplementary Table 1, Methods). To integrate the external annotation data into GPT-4o, we employed Retrieval-Augmented Generation (RAG), which improves model answers by searching external datasets and then feeding retrieved relevant information along with user input to the model.

We built RAG for GPT-4o by first constructing a search index of genetic variants on the Azure AI Search platform, using our 5 variant annotation datasets, and then connecting the search index with GPT-4o deployed at Azure OpenAI (Methods). Our search index allows for searches of variants by chromosome and position, dbSNP ID, gene, and/or condition. Other annotation fields are also searchable, although their accessibility may vary depending on the analyzer and tokenization process. The retrieved variants are ranked based on the similarity between the user input and the variant annotations. This search process is facilitated by GPT- 4o language comprehension ability, which interprets user input, generates intents from user conversations, searches and retrieves relevant information from the search index, and generates enhanced answers using the retrieved data.

We successfully incorporated 190 million variant annotations into the GPT-4o model. To evaluate information accuracy in the RAG enhanced GPT-4o output, we created test sets of 100 variants randomly selected from each dataset. We queried variant annotations using the chromosome and position of each variant and checked the output accuracy of all annotation fields, including gene, condition, IDs, allele frequencies, molecular consequences, etc. The model achieved 100% accuracy for all annotation fields in the test sets (Figure 2a). The original GPT-4o model tends to provide general information about variants, such as the name of potentially relevant datasets, which is not particularly helpful for variant interpretation and sometimes the answer includes incorrect detailed information for specific input variants (Figure 2b). After integrating annotations by RAG into GPT-4o, the model was able to deliver highly accurate and specific annotations for the user-queried variants (Figure 2c). Beyond high accuracy, the enhanced model also improved variant interpretation through its powerful reasoning capabilities, such as summarizing variant annotations (Figure 2c), suggesting disease risk based on variant allele frequency in specific populations, or generating programming scripts to visualize variants in chromosomes using the IGV tool (18). More powerful use cases and applications of this model will be discussed in Section 3.5.

**Fig. 2.**
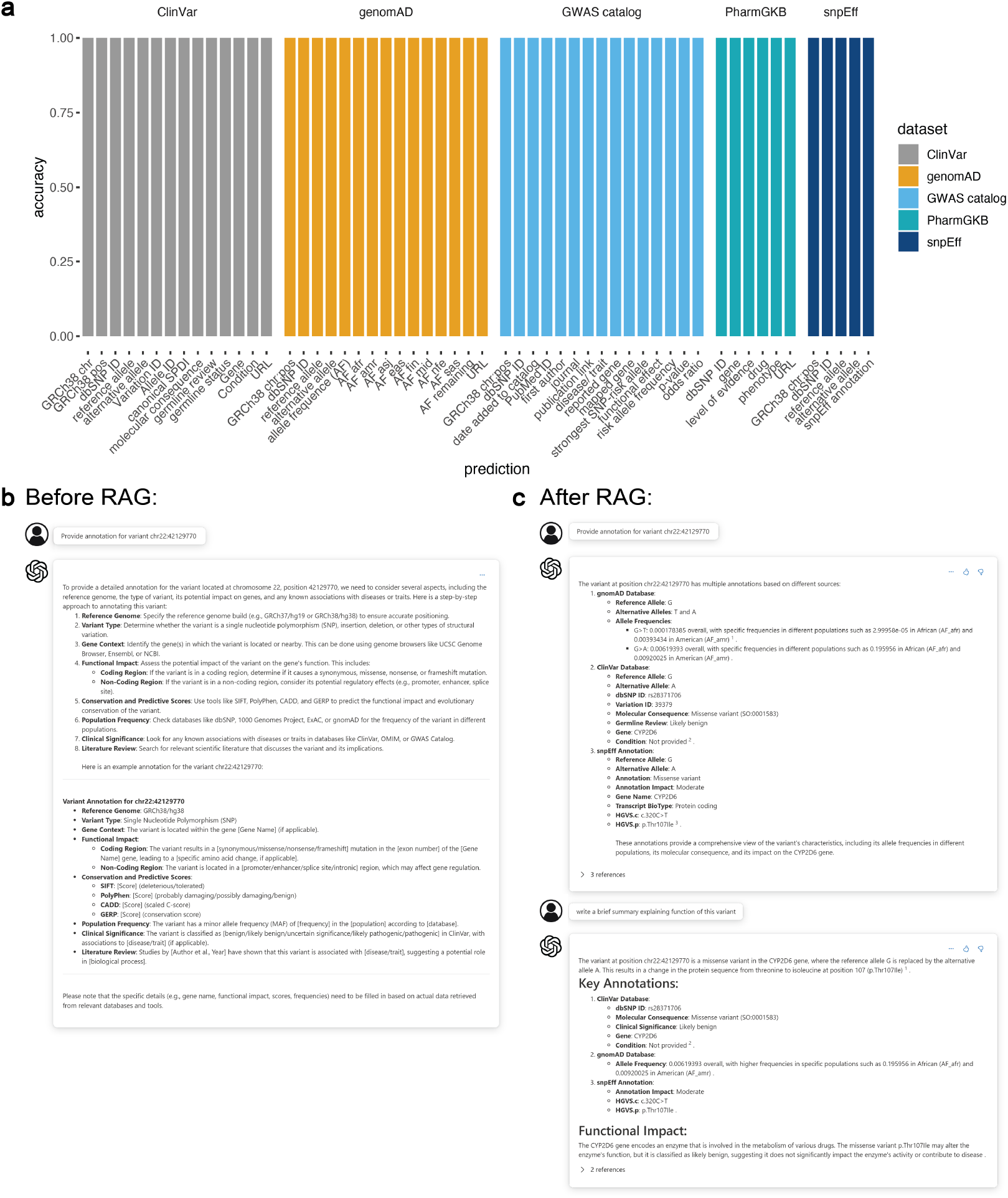
Integrating variant annotation data into GPT-4o through RAG allows the model to provide accurate annotations and enhanced interpretation. (a) Performance of RAG enhanced GPT-4o model in predicting all variant annotation fields (from 5 datasets) based on input variants. With RAG, GPT-4o achieved 100% accuracy for all annotation fields in the test sets. (b) Example dialogue with GPT-4o on variant annotations before using RAG. GPT-4o provided only general information. (c) Example dialogue with GPT-4o on variant annotations after using RAG. GPT-4o provided accurate annotations and enhanced interpretations.

### 3.3 Fine-tuning GPT-4 on variant annotation prediction

We also investigated the effectiveness of variant annotation knowledge injection into LLMs through fine-tuning. We created 3,000 training prompts and 1,000 validation prompts using variants and annotations randomly selected from the ClinVar dataset (Figure 3a). In both training and validation prompts, we configured the system message to guide the model’s focus on genomics annotation. We formatted the user input as a natural language conversation containing the chromosome and position of a variant (e.g., chr16:14555693) and instructed the model to output variant annotations. The desired output, encompassing all 13 annotation fields extracted from ClinVar, was also provided in prompts (Methods). The 100 test prompts contained the same system message and user query format as the training prompts. Our 100-variant test set was a subset of the training data, designed to evaluate the model’s ability to memorize and accurately reproduce the trained information. Despite testing multiple input formats and repetition strategies for fine-tuning GPT-4, the average output accuracy for each annotation field remained around 0.2 (Supplementary Figure 1).

**Fig. 3.**
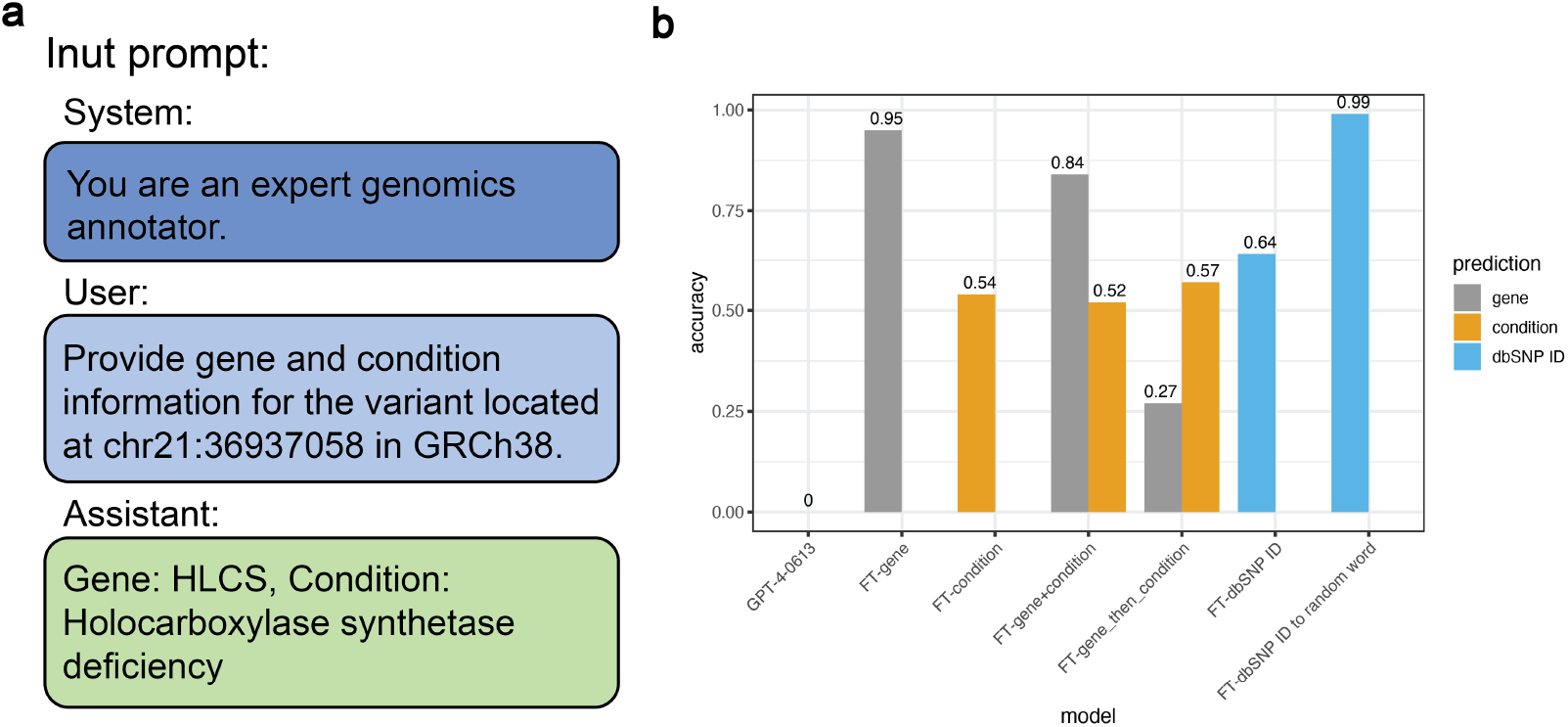
Integrating variant annotations into GPT-4 through fine-tuning. (a) Example prompt used for fine-tuning GPT-4. We provided a system message to guide the model and user messages for input variant and instruction and trained the model to predict variant annotations (assistant message). (b) Performance comparison between the base GPT-4 model and fine-tuned GPT-4 models in predicting genes, conditions, or dbSNP IDs.

To improve fine-tuned model’s performance, we adjusted our approach by fine-tuning GPT-4 to learn a single annotation field at a time (Figure 3b). We selected variants from 300 genes, with 10 variants per gene, and fine-tuned the model specifically on predicting the gene field, achieving an accuracy of 0.95. When we fine-tuned a GPT-4 model to predict the condition field using the same dataset, the accuracy dropped to 0.54. This lower accuracy was due to the overrepresentation of the ‘not provided’ category in conditions, leading the model to learn to predict ‘not provided’ for all cases. For dbSNP ID prediction (e.g., rs28371706), we fine-tuned the model on dbSNP IDs for 300 variants with 10 repeats for each. The initial accuracy was 0.64. By converting each ID into a word represented by a single token to improve the tokenization process, we managed to increase the prediction accuracy to 0.99. When we expanded the task to predict both the gene and condition fields simultaneously, the accuracy decreased to 0.84 for the gene field and 0.52 for the condition field. Continuous fine-tuning of the condition field on the already fine-tuned gene model did not result in any further improvement in prediction accuracy.

This led us to conclude that fine-tuning GPT-4 on variant annotation data can improve performance in certain annotation fields, such as gene and dbSNP ID, although the overall accuracy across multiple fields remains suboptimal. We found that as we incorporated more diverse information and dealt with less frequently occurring data, the model faced increasing challenges in learning effectively through fine-tuning.

### 3.4 Comparison between RAG and fine-tuning

Our results indicated that RAG outperforms fine-tuning when injecting knowledge into LLMs in terms of data volume, accuracy, cost, time efficiency, and flexibility. Using RAG, we successfully integrated 190 million variant annotations. In contrast, fine-tuning struggled to add 13 annotation fields of 3,000 variants to the LLM. As for accuracy, RAG ensured 100% accuracy relying on retrieving correct documents. Fine-tuning, however, found it challenging to achieve such high accuracy levels. The primary cost for RAG lies in creating and storing the search index, which depends on the indexing platform. In our case, we used the Azure AI Search Standard S1 price tier, costing approximately 250*permonth*(*Aug*2024).*Fine − tuning, ontheotherhand, costs*25 per million input tokens. Fine-tuning 3,000 variant annotations for three epochs billed approximately 1 million training tokens. Scaling this to 190 million annotations would be significantly more expensive than using RAG. As for time efficiency, the time required to build a RAG index depends on the search method employed. We used keyword search, which took approximately 30 hours to create the search index for all 190 million annotations. In comparison, fine-tuning 3,000 annotations for three epochs took about 4 hours. Extending this process to 190 million variants, even assuming a linear increase rate, would require a substantial amount of time. Another advantage of RAG is its flexibility to plug into any LLM, allowing users to choose their preferred base model. It can also be easily added to new models when model’s updates occur. Fine-tuning, however, relies on a specific model and cannot be easily migrated.

### 3.5 Use cases of RAG enhanced GPT-4o model

The potential use cases of our RAG GPT-4o model extend far beyond providing highly accurate annotations. To illustrate our model’s capabilities, we presented an example use case simulating a disease diagnosis scenario. In this scenario, given the symptoms of a patient and their variant list reported by genetic tests, the model inferred the disease and identified causal variants (Figure 4).

**Fig. 4.**
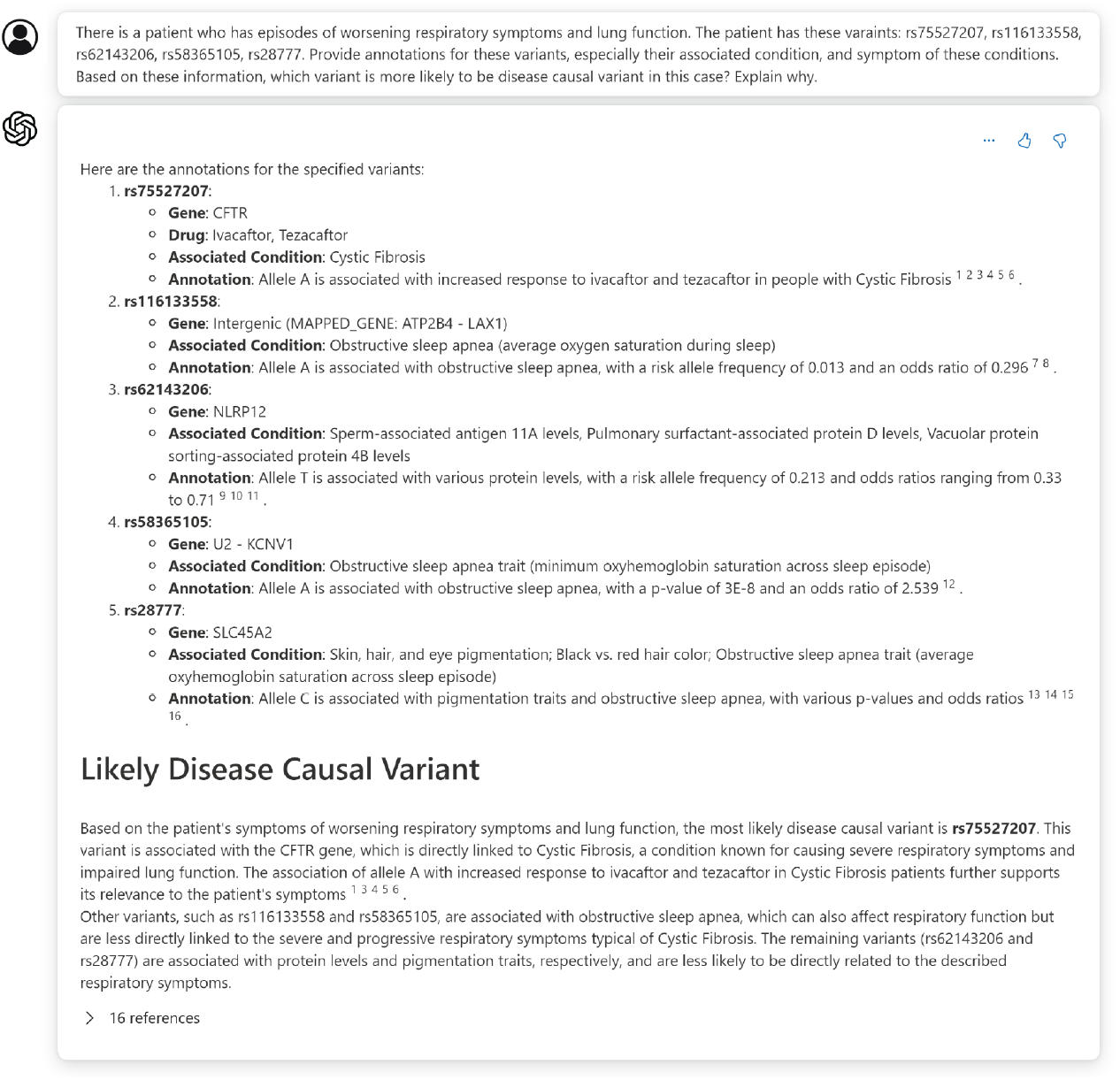
Example dialogue of a disease-causing variant identification scenario using the RAG enhanced GPT-4o model. In this example, we provided the model with disease symptoms and a list of variants. Our model successfully provided accurate variant annotations and identified both the associated disease and the disease-causing variant from the list.

We input a typical symptom of cystic fibrosis, episodes of worsening respiratory symptoms and lung function, along with a variant list containing a cystic fibrosis causal variant (rs75527207), and 4 random variants unrelated to cystic fibrosis (rs116133558, rs62143206, rs58365105, rs28777). We then queried the model about potential disease-causing variants in this case using a natural language conversation.

With variant annotations provided by the dataset we injected through RAG, the model accurately provided the related disease and annotations for the input variants. Beyond this initial step, the model identified the symptoms of the diseases, compared the similarities between disease symptoms and the input symptoms using its knowledge and powerful logic and reasoning capabilities, and generated an informative answer, with a detailed explanation of why it concludes that the variant is more likely to be causal. This significantly reduces the workload involved in disease diagnosis, eliminating the need to manually identify and search related variant databases, gather symptoms, and generate hypotheses.

## 4. Discussion

We have significantly improved the performance of the GPT- 4o model in genomics by successfully integrating 190 million variant annotations derived from 5 datasets that cover most variants and their annotations. Our model enables users to query specific variants using natural language conversation, by chromosome and position, dbSNP ID, gene, condition, or other annotation fields. Users receive accurate annotations and informative interpretations, supported by GPT-4o’s powerful reasoning and conversation comprehension capabilities. This improvement makes the variant interpretation process more efficient and accessible for healthcare providers and researchers.

We also explored the use of small language models (SLMs), such as Phi-3 (19), for incorporating variant annotations. Despite having fewer parameters and less training data compared to LLMs, SLMs are still highly capable in many tasks. These models require fewer resources to run and can be deployed to a phone to process data locally. This characteristic makes SLMs particularly valuable for handling private genomics data, as they provide an option to enhance variant interpretation while simultaneously ensuring the security and privacy of sensitive information. We successfully incorporated 1,000 variant annotations from ClinVar into the Phi-3 model using Semantic Kernel (v.1.16.1) (20) significantly improving the quality of variant annotation query responses. Further exploration is needed to integrate more variants and continue to improve answer quality.

Our study has demonstrated the capabilities of our RAG GPT-4o model, highlighting its potential in various scenarios. However, it is important to acknowledge the existing limitations of our model. One such constraint is the lack of comprehensive other background genomics knowledge in GPT-4o. For instance, the model did not understand that high allele frequency variants tend to be benign and unlikely to be disease-causing variants, which affects answer quality for related questions. Additionally, the search method we employed in RAG is keyword-based, which limits the retrieval of relevant documents to user input keywords. Exploring vector search techniques could accommodate a wider range of user input scenarios, retrieve more relevant documents, and reduce response latency. Our model can be further enhanced by incorporating multimodality genomics data and improving global question-answering capabilities through techniques such as vector search or GraphRAG (21).

As a pioneering study integrating genomics data with natural conversation LLMs, our work paves the way for the development of more comprehensive and helpful genomics AI systems to support disease diagnosis and facilitate research discovery in the future.

## 6. Disclaimer

While we strive to ensure the accuracy and reliability of the information generated by this large language model, we cannot guarantee its completeness or correctness. Users should independently verify any information and consult with experts in the respective field to obtain accurate and up-to-date advice.

## 7. Acknowledgement

We thank Jeremiah (Miah) Wander (Microsoft Health Future) for discussions, sharing ideas about evaluation and potential use cases of our LLM models.

