## Supplemental Table 1 and Supplemental Figure 1 for "Boosting GPT Models for Genomics Analysis: Generating Trusted Genetic Variant Annotations and Interpretations through RAG and fine-tuning"

**Supplementary Table 1**: Overview of variant annotation datasets integrated into GPT models through RAG or fine-tuning.

| Dataset/ Tool | No. Variants | Selected Annotation Fields (one example variant) |
|---|---|---|
| ClinVar | 2,897,556 | GRCh38 chr: chr1<br>GRCh38 pos: 69134<br>reference allele: A<br>alternative allele: G<br>dbSNP ID: na<br>Variation ID: 2205837<br>Allele ID: 2193183<br>canonical SPDI: NC_000001.11:g.69134A>G<br>molecular consequence: SO:0001583\|missense_variant<br>germline review: Likely benign<br>germline status: criteria provided, single submitter<br>Gene: OR4F5<br>Condition: not specified<br>source: ClinVar<br>ClinVar URL:<br>https://www.ncbi.nlm.nih.gov/clinvar/variation/2205837/ |
| gnomAD | 183,717,261 | GRCh38 chr:pos: chr1:12948<br>dbSNP ID: rs1199063229<br>reference allele: T<br>alternative allele: C<br>allele frequence (AF): 0.000367962<br>AF afr: 0.00523088<br>AF amr: 0.000373227<br>AF asj: 0.000291545<br>AF eas: 0.000439947<br>AF fin: 0<br>AF mid: 0.00174825<br>AF nfe: 0.000132535<br>AF sas: 8.98796e-05<br>AF remaining: 0.000959003<br>source: gnomAD<br>gnomAD URL: https://gnomad.broadinstitute.org/variant/1-12948-T-C?dataset=gnomad_r4 |
| SnpEff | 2,897,556 | GRCh38 chr:pos: chr1:69134<br>dbSNP ID: na<br>reference allele: A<br>alternative allele: G<br>allele: G<br>annotation: missense variant |

| | | |
|---|---|---|
| | | annotation Impact: MODERATE<br>gene name: OR4F5<br>gene ID: ENSG00000186092.7<br>feature type: transcript<br>feature ID: ENST00000641515.2<br>transcript BioType: protein coding<br>rank: 3/3<br>HGVS.c: c.107A>G<br>HGVS.p: p.Glu36Gly<br>cDNA.pos/cDNA.length: 167/2618<br>CDS.pos/CDS.length: 107/981<br>AA.pos/AA.length: 36/326<br>Distance:<br>ERRORS/WARNINGS/INFO:<br>source: snpEff |
| GWAS Catalog | 625,113 | GRCh38 chr:pos: chr3:85356351<br>dbSNP ID: rs9822731<br>date added to catalog: 2019-03-18<br>PubMed ID: 30643258<br>first author: Karlsson Linner R<br>journal: Nat Genet<br>publication link:<br>https://www.ncbi.nlm.nih.gov/pubmed/30643258<br>disease/ trait: Alcohol consumption (drinks per week)<br>reported gene: CADM2<br>mapped gene: CADM2<br>strongest SNP-risk allele: rs9822731-T<br>functional effect: intron variant<br>risk allele frequency: 0.7753<br>p-value: 4E-15<br>odds ratio: 0.021035347<br>source: GWAS Catalog |
| pharmGKB | 41,287 | dbSNP ID: rs75527207<br>gene: CFTR<br>level of evidence:1A<br>drug: ivacaftor<br>phenotype: Cystic Fibrosis<br>source: pharmGKB<br>pharmGKB URL:<br>https://www.pharmgkb.org/clinicalAnnotation/981755803 |

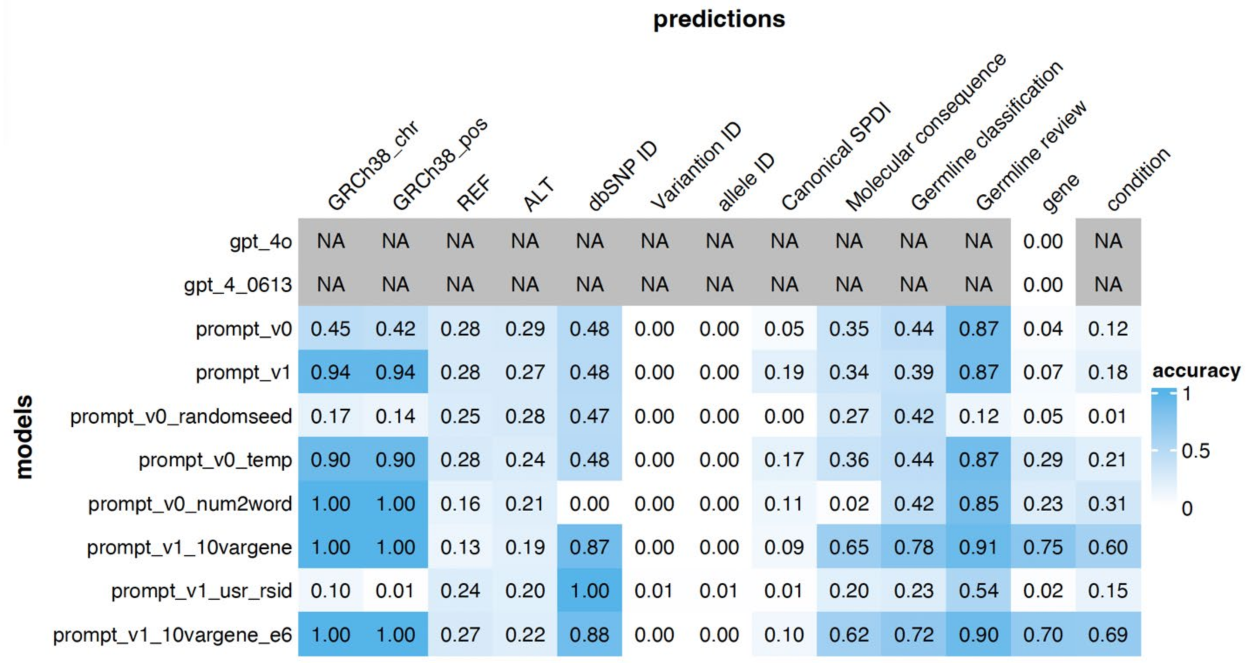


**Supplementary figure1**. Fine-tuning GPT-4 model for predicting 13 annotation fields simultaneously. The performance of each model is measured by the accuracy of predicting the exact match of each annotation field. The base GPT-4o and GPT-4 models are unable to correctly predict genes from user provided variants. We fine-tuned the models using diverse input data formats, including: 1) prompt_v0: used the same format as described in the Methods section; 2) prompt_v1: added { } around each annotation field to enhance segmentation; 3) randomseed: tested the model with a different random seed; 4) temp: tests model with higher temperature setting; 5) num2word: encoded numerical IDs (dbSNP ID, Variation ID, and allele ID) as random words to improve tokenization; 6) 10vargene: included 10 variants per gene in the training dataset, 7) usr_rsid: changed user input variants from the chr:pos format to the dbSNP ID; 8) 10vargene_e6: included 10 variants per gene in the training dataset and trained model for an addition 3 epochs, resulting in 6 epochs in total.
